# Evolutionary trade-offs at the Arabidopsis *WRR4A* resistance locus underpin alternate *Albugo* candida recognition specificities

**DOI:** 10.1101/2021.03.29.437434

**Authors:** Baptiste Castel, Sebastian Fairhead, Oliver J. Furzer, Amey Redkar, Shanshan Wang, Volkan Cevik, Eric B. Holub, Jonathan D. G. Jones

**Affiliations:** The Sainsbury Laboratory, University of East Anglia, Norwich Research Park, NR4 7UH, Norwich, United Kingdom; Department of Biological Sciences, National University of Singapore, Singapore 117558; Warwick Crop Centre, School of Life Sciences, University of Warwick, CV35 9EF, Wellesbourne, United Kingdom; Department of Biology, University of North Carolina, Chapel Hill, NC 27599, USA; Department of Genetics, University of Cordoba, 14071 Cordoba, Spain; The Milner Centre for Evolution, Department of Biology and Biochemistry, University of Bath, BA2 7AY Bath, United Kingdom

**Keywords:** immunity, resistance gene, NLR, natural variation, evolution, effector recognition, crop protection, Arabidopsis thaliana, camelina

## Abstract

The oomycete *Albugo candida* causes white rust of Brassicaceae, including vegetable and oilseed crops, and wild relatives such as *Arabidopsis thaliana*. Novel *White Rust Resistance* (*WRR*)-genes from Arabidopsis enable new insights into plant/parasite co-evolution. *WRR4A* from Arabidopsis accession Col-0 provides resistance to many but not all white rust races, and encodes a nucleotide-binding (NB), leucine-rich repeat (LRR) (NLR) immune receptor protein. Col-0 *WRR4A* resistance is broken by a Col-0-virulent isolate of *A. candida* race 4 (AcEx1). We identified an allele of *WRR4A* in Arabidopsis accession Oy-0 and other accessions that confers full resistance to AcEx1. *WRR4A*^*Oy-0*^ carries a C-terminal extension required for recognition of AcEx1, but reduces recognition of several effectors recognized by the *WRR4A*^*Col-0*^ allele. *WRR4A*^*Oy-0*^ confers full resistance to AcEx1 when expressed as a transgene in the oilseed crop *Camelina sativa*.

**Significance:** A C-terminal extension in an allele of the Arabidopsis resistance-protein WRR4A changes effector recognition specificity, enabling the *WRR4A*^*Oy-0*^ allele to confer immunity to *Albugo candida* races that overcome the *WRR4A*^*Col-0*^ allele. This resistance can be transferred to the oil-producing crop *Camelina sativa*.

**Graphical abstract:** 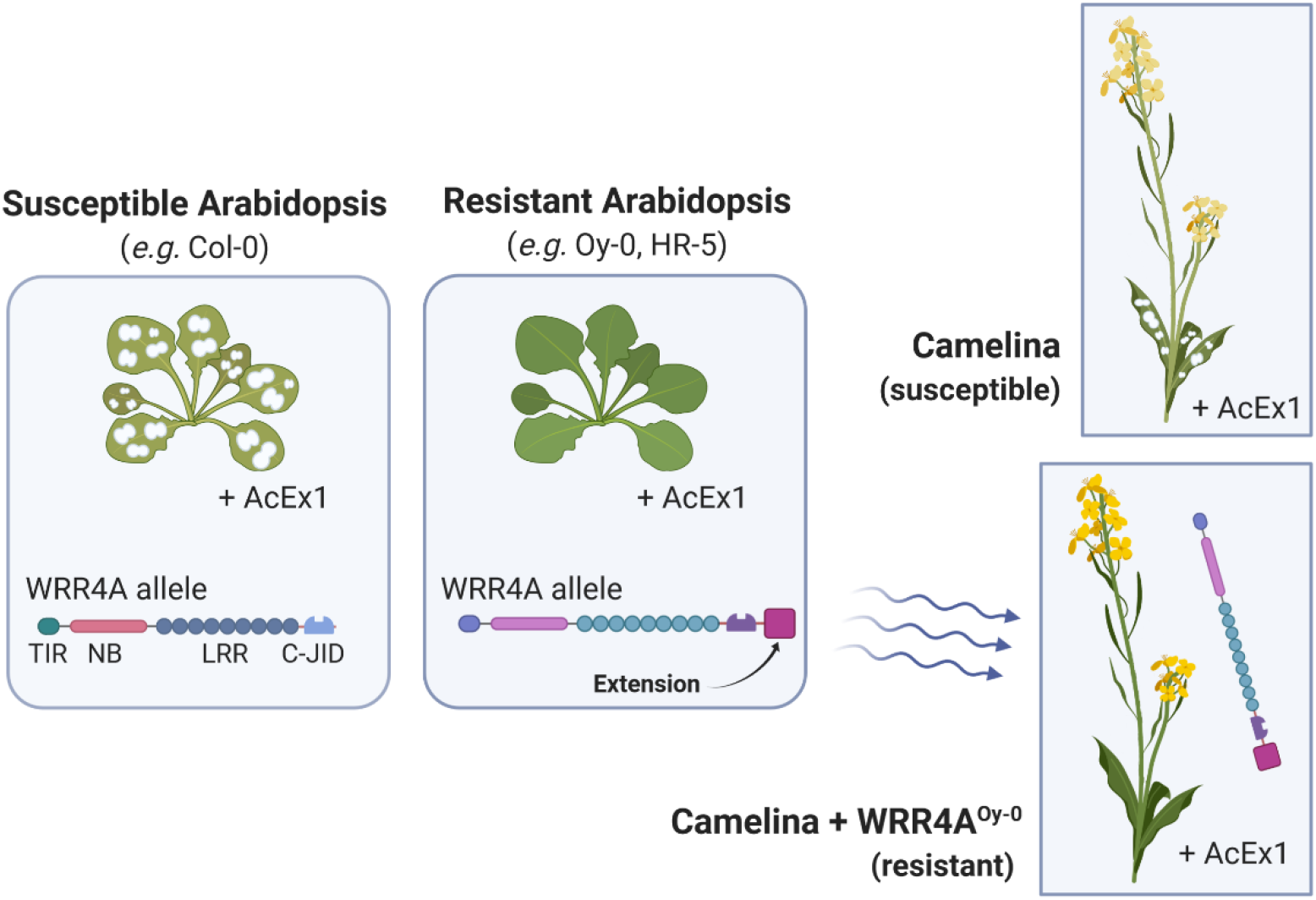

## Introduction

Plants have evolved powerful defense mechanisms that can arrest attempted colonization by microbial pathogens. Timely defense activation requires perception of pathogen-derived molecules by cell-surface pattern-recognition receptors (PRRs) and intracellular Nucleotide-binding (NB), Leucine-rich repeat (LRR), or NLR, immune receptors (Jones and Dangl, 2006). Extensive NLR genetic diversity within plant populations is associated with robustness of NLR-mediated immunity (Baggs *et al*., 2017), and plant NLR sequences reveal diversifying selection on NLR genes compared to other genes (Kuang *et al*., 2004; Monteiro and Nishimura, 2018; Meyers *et al*., 1998). To investigate NLR diversity, next-generation sequencing technologies were combined with sequence capture to develop *Resistance (R)*-gene enrichment sequencing (RenSeq) (Jupe *et al*., 2013). This method has shed new light on NLR repertoires in several plant genomes including tomato, potato and wheat (Andolfo *et al*., 2014; Witek *et al*., 2016; Steuernagel *et al*., 2016). A comparison of 64 *Arabidopsis thaliana* (Arabidopsis) accessions using RenSeq documented NLR sequence diversity within a single species, revealing the Arabidopsis “pan-NLRome” (Van de Weyer *et al*., 2019). Each Arabidopsis accession contains 150-200 NLR-encoding genes. About 60% are found in clusters (within 200 kb from each other) that show copy number variation (Lee and Chae, 2020). From all the NLRs of the 64 accessions, 10% are singletons and the rest are distributed between 464 orthogroups. Each accession contains a unique subset comprising, on average, 25% of the orthogroups.

NLRs vary in their intramolecular architecture. Plant NLR proteins usually display either a “Toll, Interleukin-1, *R*-gene” (TIR), a “Coiled-Coil” (CC) or “Resistance to Powdery mildew 8” (RPW8) N-terminal domain, a central NB domain and a C-terminal LRR domain. Some NLRs also comprise a post-LRR (PL) domain. For example, RRS1 is an Arabidopsis TIR-NLR with PL WRKY domain required to detect the effectors AvrRps4 (from the bacterium *Pseudomonas syringae*) and PopP2 (from the bacterium *Ralstonia solanacearum*). The integrated WRKY is called a decoy as it mimics the authentic AvrRps4 and PopP2 effector targets (Sarris *et al*., 2015; Le Roux *et al*., 2015). Several other integrated decoy domains have been described (Cesari, 2017). RPP1 and Roq1, two TIR-NLRs from Arabidopsis and *Nicotiana benthamiana* respectively, form tetrameric resistosomes upon activation. In this structure, a PL C-terminal jelly-roll/Ig-like domain (C-JID) physically binds the cognate effector, along with the LRR domain (Ma *et al*., 2020; Martin *et al*., 2020). Analysis of NLR integrated domains can potentially reveal novel effector targets (Kroj *et al*., 2016).

*A. candida* causes white blister rust in Brassicaceae and serious annual yield losses in brassica crops such as oilseed mustard (*Brassica juncea*) in India (Gupta *et al*., 2018). It comprises several host-specific subclades, which includes race 2 from *B. juncea*, race 7 from *Brassica rapa*, race 9 from *Brassica oleracea* and race 4 from crop relatives (*e*.*g*., *Capsella bursa-pastoris, Arabidopsis spp*. and *Camelina sativa*) (Jouet *et al*., 2018; Pound and Williams, 1963). These have been proposed to evolve by rare recombination events that occurred between the races, followed by clonal propagation on susceptible hosts (McMullan *et al*., 2015). The Arabidopsis Columbia (Col-0) allele of *WRR4A* can confer resistance to isolates of all four races (Borhan *et al*., 2010; Borhan *et al*., 2008). The allele encodes a canonical TIR-NLR and belongs to an orthogroup of three genes in Col-0 at the same locus. The accession Ws-2 (susceptible to *A. candida* race 4) lacks *WRR4A* but contains the two other paralogs, illustrating intra-species copy number variation within clusters. Interestingly, one of these paralogs, *WRR4B*, also confers resistance to the Ac2V isolate of race 2 (Cevik *et al*., 2019). In addition, the CC-NLR-encoding *BjuWRR1*, which confers resistance to several *candida* isolates collected on *B. juncea*, was mapped and cloned from the European accession of *B. juncea* Donskaja-IV (Arora *et al*., 2019).

Several Col-0-virulent isolates of *A. candida* race 4 have been collected from naturally infected Arabidopsis plants. They were used subsequently to identify an alternative source of broad-spectrum white rust resistance. One was found in Arabidopsis accession Oy-0 that mapped to the *WRR4* locus (Fairhead, 2016). Interestingly, one of these isolates (AcEx1), as well as the related white rust pathogen *Albugo laibachii*, were shown to suppress non-host resistance in Arabidopsis to the potato late blight pathogen *Phytophthora infestans* (Prince *et al*., 2017; Belhaj *et al*., 2017). Thus, we set out to clone the gene conferring AcEx1 resistance in Oy-0, and characterise the corresponding pathogen effector(s).

AcEx1 is also virulent in *Camelina sativa*, which is an emerging oilseed crop and has been engineered to provide an alternative source of fish-oil-derived long chain omega-3 polyunsaturated fatty acids (LC-PUFAs). Algal-derived genes were expressed in the seed to produce eicosapentaenoic acid (EPA) and docosahexaenoic acid (DHA) as an improved LC-PUFA source for fish feed or as a human nutritional supplement (Ruiz-Lopez *et al*., 2014; Petrie *et al*., 2014). EPA and DHA, recognised for their health benefits, are mainly sourced from oily fish, and are acquired from feeding on algae-consuming plankton. In salmon farming, the major source for of omega-3 oil derives from oceanic fish. Industry collects 750,000 metric tons of fish oil every year, raising sustainability concerns (Napier *et al*., 2015). Transgenic camelina oil is equivalent to fish oil for salmon feeding and for human health benefits (Betancor *et al*., 2018; West *et al*., 2019). Despite challenges to distribute a product derived from a genetically modified crop (Napier *et al*., 2019), an increase in camelina cultivation can be expected in the near future. Fields of *C. sativa* will be exposed to *A. candida* and early identification of *R*-genes will enable crop protection.

In this study we identified two alleles of *WRR4A* conferring full resistance to AcEx1 from Arabidopsis accessions Oy-0 and HR-5. They both encode proteins with a C-terminal extension compared to the Col-0 *WRR4A* allele. This extension enables recognition of at least one effector from AcEx1. We propose that *WRR4A*^Oy-0^ is the ancestral state, and that in the absence of AcEx1 selective pressure, an early stop codon in *WRR4A* generated the Col-0-like allele, enabling more robust recognition of other *A. candida* races while losing recognition of AcEx1. Finally, we successfully transferred *WRR4A*^*Oy-0*^-mediated resistance to AcEx1 from Oy-0 into *Camelina sativa*.

## Results

### *Resistance to AcEx1 is explained by* WRR4A *alleles of HR-5 and Oy-0*

AcEx1 growth on Col-0 results in chlorosis that is not seen in the fully susceptible accession Ws-2 (**Figure 1a**). Since *WRR4A* confers resistance to all other *A. candida* races tested and Ws-2 lacks *WRR4A*, we tested if the chlorotic response could be explained by *WRR4A*, by testing a Col-0_*wrr4a-6* mutant, and found that it shows green susceptibility to AcEx1. We also tested Ws-2 transgenic lines carrying *WRR4A* from Col-0 and observed chlorotic susceptibility (**Figure 1a**). Thus, *WRR4A* from Col-0 weakly recognises AcEx1 and provides partial resistance. However, AcEx1 is still able to complete its life cycle on Col-0.

**Figure 1:**
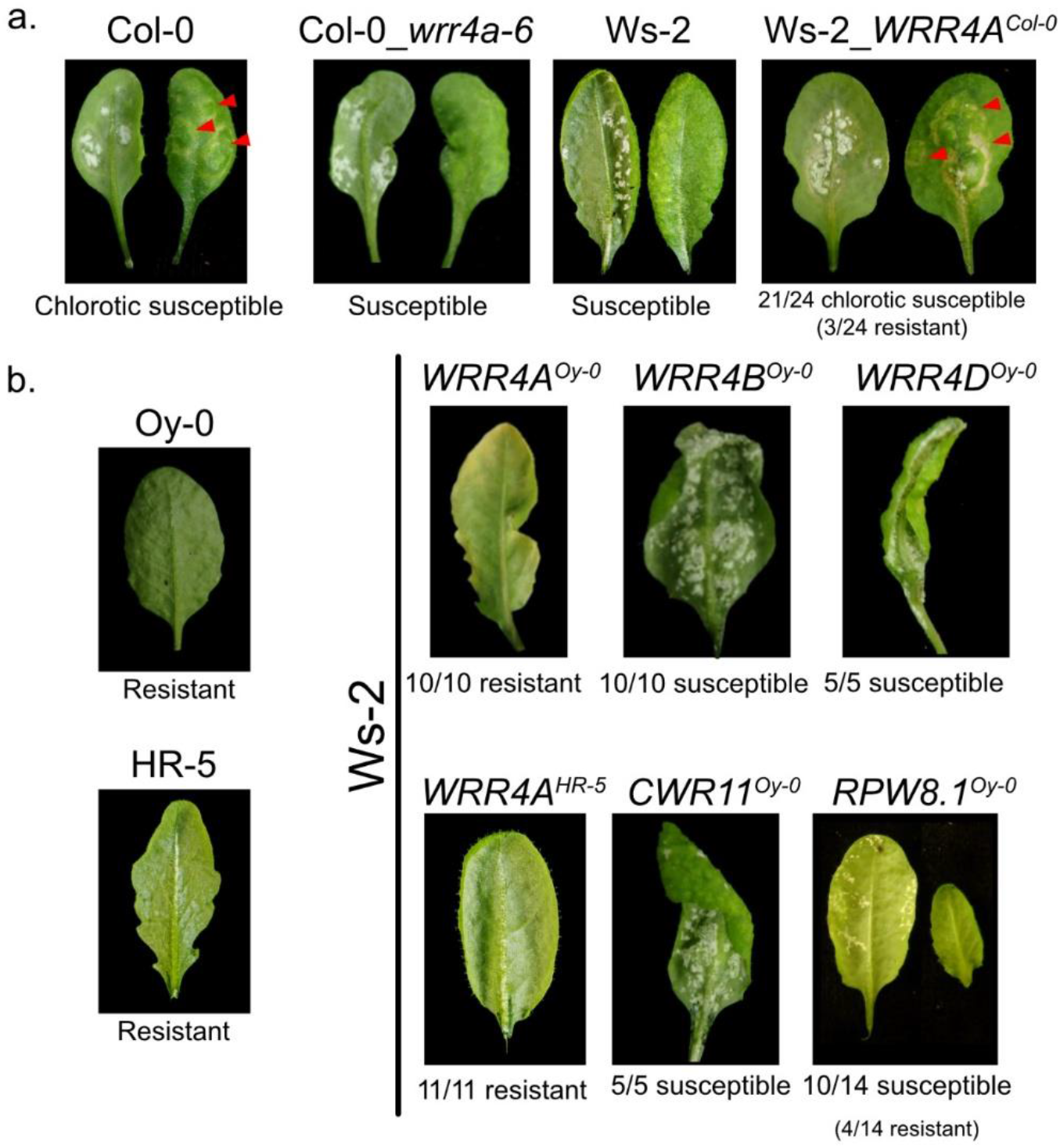
Oy-0 and HR-5 alleles of WRR4A confers full resistance to AcEx1. 5-week old plants were sprayed inoculated with AcEx1. Plants were phenotyped 14 days after inoculation. **a**. Col-0 and Ws-2_*WRR4A*^*Col-0*^ present a chlorotic susceptible, indicated by red arrows; Col-0_*wrr4a-6* and Ws-2 do not. **b**. Oy-0 and HR-5 are fully resistant, as well as Ws-2_*WRR4A*^*Oy-0*^ and Ws-2_*WRR4A*^*HR-5*^. Ws-2 expressing *WRR4B*^*Oy-0*^, *WRR4D*^*Oy-0*^ or *CWR11*^*Oy-0*^ are fully susceptible. Ws-2 lines expressing *RPW8*.*1*^*Oy-0*^ are generally susceptible, 4/14 showed dwarphism and resistance. Numbers indicate the number lines showing similar phenotype out of the number of plants tested.

In a search for more robust sources of AcEx1 resistance, we tested 283 Arabidopsis accessions (**Table S2**). We identified 57 (20.1%) fully resistant lines, including Oy-0 and HR-We phenotyped 278 Recombinant Inbred Lines (RILs) between Oy-0 (resistant) and Col-0 (susceptible) and conducted a quantitative trait locus (QTL) analysis that revealed one major QTL on chromosome 1 and two minor QTLs on chromosomes 3 and 5 (**Figure S1a**). All loci contribute to resistance, with a predominant contribution of the QTL on chromosome 1 (see Figure 3.7 of Fairhead, 2016). We did not investigate the minor QTL on chromosome 5. Fine mapping on chromosome 1 and 3 QTLs refined the QTL boundaries (**Figure S2 and S3**, see Experimental Procedures). Based on sequence identity between the QTL in Col-0 and in an Oy-0 RenSeq dataset (Van de Weyer *et al*., 2019), we identified four NLRs associated with the QTLs in Oy-0: three TIR-NLR paralogs on chromosome 1 (*WRR4A, WRR4B* and one absent in Col-0 that we called *WRR4D*) and a CC-NLR absent in Col-0 on chromosome 3 (that we called *Candidate to be WRR11, CWR11*) (**Figure S1B-C**).

We expressed these genes, with their own promoters and terminators, in the fully susceptible accession Ws-2. Only *WRR4A*^*Oy-0*^ conferred full resistance (**Figure 1b**). *CWR11*, the only NLR from the *WRR11* locus, does not confer AcEx1 resistance. The *WRR11* locus is orthologous to the position 17.283-18.535 Mb on chromosome 3 of Col-0 (**Figure S3**). The RPW8 alleles of Col-0 are encoded between positions 18.722-18.734 Mb. Oy-0 carries 15 copies of *RPW8* (Van de Weyer *et al*., 2019), so it could be that some copies recombined on the other side of the *WRR11* locus. We cloned one homolog of RPW8.1 from Oy-0 and expressed it with its own promoter and terminator in Ws-2 (**Figure 1**). Most transgenic lines were susceptible to AcEx1 but four lines were resistant. These lines were also smaller than the other lines, suggesting an autoimmune phenotype. Since most transgenic lines are susceptible and ectopic expression of RPW8 is known to result in autoimmunity (Xiao *et al*., 2003), we did not further investigate the role of RPW8^Oy-0^ paralogs in AcEx1 resistance. The gene underlying *WRR11* locus resistance remains unknown.

We conducted a bulk segregant analysis using an F2 population between HR-5 (resistant) and Ws-2 (susceptible). RenSeq on bulked F2 susceptible segregants revealed a single locus on chromosome 1, that maps to the same position as the chromosome 1 QTL in Oy-0 (**Figure S4A**). Since WRR4A^Oy-0^ confers resistance to AcEx1, we expressed its HR-5 ortholog, in genomic context, in the fully susceptible accession Ws-2, and found that *WRR4A*^*HR-5*^ also confers full resistance to AcEx1 (**Figure 1b**).

In conclusion, *WRR4A* from Col-0 can weakly recognise AcEx1 but does not provide full resistance. We identified two *WRR4A* alleles, in Oy-0 and HR-5, that confer complete AcEx1 resistance.

### WRR4A^*Col-0*^ *carries an early stop codon compared to WRR4A*^*Oy-0*^

To understand why the Oy-0 and HR-5 alleles of *WRR4A* confer full resistance to AcEx1, while the Col-0 allele does not, we compared the gene and protein sequences (**Figure 2**). First, we defined the cDNA sequence of *WRR4A*^*Oy-0*^. The splicing sites are identical between the two alleles. There are 46 polymorphic amino acids between Col-0, HR-5 and Oy-0. Col-0 shares 96.03% identity with Oy-0 and 96.23% with HR-5, while Oy-0 and HR-5 share 97.15% identity. *WRR4A*^*Col-0*^ carries a 156-nucleotide insertion in the first intron compared to Oy-0 and HR-5. A striking polymorphism is a TGC->TGA mutation in *WRR4A*^*Col-0*^, resulting in an early stop codon compared to *WRR4A*^*Oy-0*^ and *WRR4A*^*HR-5*^ (**Figure 2**), located 178 amino acids after the last LRR, resulting in an 89 amino acid extension in WRR4A^Oy-0^ and WRR4A^HR-5^. The nucleotide sequence for this extension is almost identical between HR-5, Oy-0 and Col-0 (two polymorphic sites). Thus, by mutating TGA to TGC in Col-0, we could engineer an allele with the extension, that we called *WRR4A*^*Col-0_LONG*^ (**Figure 3a**). By mutating TGC to TGA in Oy-0, we could engineer an Oy-0 allele without the extension, that we called *WRR4A*^*Oy-0_SHORT*^. We expressed these alleles, as well as the WT Col-0 and Oy-0 alleles, with their genomic context, in the AcEx1-compatible accession Ws-2. None of the *WRR4A*^*Col-0_LONG*^ and *WRR4A*^*Oy-0_SHORT*^ transgenic seeds germinated, suggesting autoactivity in Arabidopsis. We tried to generate Arabidopsis Col-0 lines with *WRR4A*^*Col-0-STOP*^ using CRISPR adenine base editor (see Experimental Procedures). Out of 24 transformed plants, none displayed editing activity at all. Thus, we did not generate stable *WRR4A* stop codon mutants in Arabidopsis. However, we cloned these alleles under the control of the *35S* promoter and the *Ocs* terminator for transient overexpression in *N. tabacum*.

**Figure 2:**
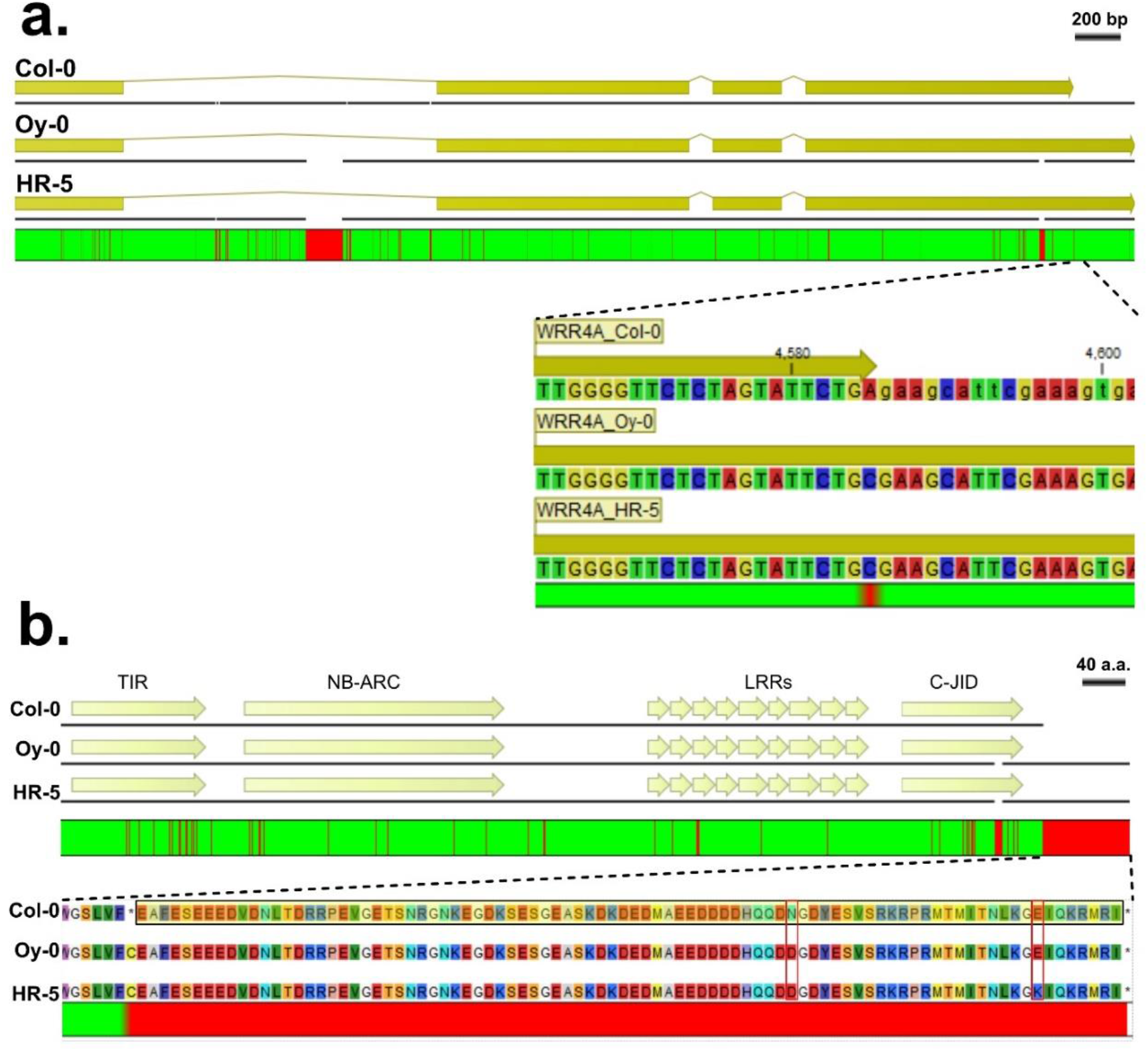
Allelic variation between Col-0 and Oy-0 alleles of WRR4A. **a**. Nucleotide sequence alignment from ATG to TGA. Plain yellow areas represent exons. Yellow lines represent introns. bp: base pair **b**. Amino acid alignment. a.a: amino acid. The C-terminal extension is framed in yellow for Col-0 to indicate that an early stop codon avoids translation of this sequence. **a.b**. Cartoons made with CLC Workbench Main. Green represents identity. Red represents polymorphism. Figures are on scale.

**Figure 3:**
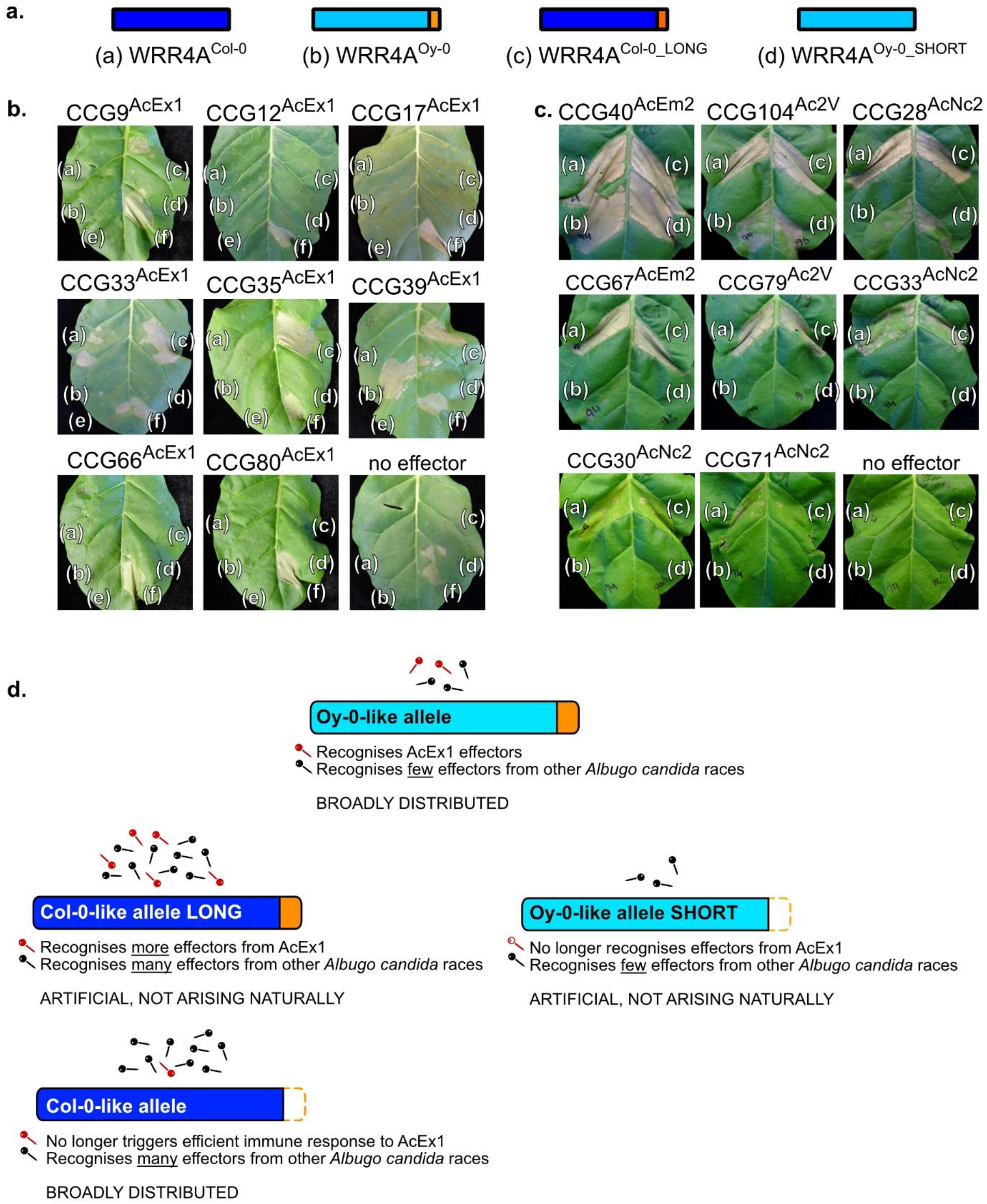
Recognition of CCG effectors by WT and stop codon mutant alleles of *WRR4A*. CCG effector candidates were transiently expressed in 4-week old *N. tabacum* leaves, under the control of the 35S promoter and Ocs terminator, alone or with WT or mutant alleles of *WRR4A*. Leaves were infiltrated with *Agrobacterium tumefaciens* strain GV3101 in infiltration buffer at OD_600_ = 0.4. Pictures were taken at 4 dpi. **a**. cartoon of the WRR4A alleles: (a) Col-0 WT, (b) Oy-0 WT, (c) Col-0 with TGA-TGC mutation, causing an Oy-0 like C-terminal extension, (d) Oy-0 with a TGC-TGA mutation causing a truncation of the C-terminal extension. **b**. AcEx1 CCG effector candidates alone (e) or with one of the four WRR4A alleles as shown in Fig 3a (a), (b), (c), (d). MLA7 CC domain was used as an HR positive control (f). **c**. Seven CCGs from other races of *Albugo candida* known to be recognized by Col-0 allele of WRR4A were tested with the three others WRR4A alleles. Co-expression of CCG with their cognate R-protein WRR4A Col-0 was used as a positive control (a). Expression of the WRR4A alleles without effector was used as negative controls (bottom right picture). **d**. Summary of the CCG recognition by WRR4A alleles. Red nails: AcEx1 effectors, black nails: CCGs from other *A. candida* races.

Since many TIR-NLRs carry a PL C-JID, we conducted a Hidden Markov Model (HMM) search for C-JID and found one in WRR4A (www.ebi.ac.uk/Tools/hmmer/search/hmmsearch on *Arabidopsis thaliana* using C-JIC HMM previously reported (Ma *et al*., 2020), e-value = 5.7e-14). This C-JID is present in both Oy-0 and Col-0 alleles (**Figure 2b**). The C-terminal extension in WRR4A^Oy-0^ relative to WRR4A^Col-0^ does not show homology with known protein domains.

### *Extension in* WRR4A *confers specific recognition of AcEx1 candidate effectors*

We tried to identify AcEx1 effectors specifically recognised by WRR4A^Oy-0^. We tested for hypersensitive response (HR), as typical marker of NLR activation, upon transient WRR4A^Oy-0^ expression along with AcEx1 effector candidates in *N. tabacum* leaves. Secreted CxxCxxxxxG (or CCGs) proteins are expanded in the genomes of *Albugo* species and are effector candidates (Kemen *et al*., 2011, Furzer *et al*., 2021). We identified 55 CCGs in the AcEx1 genome (Jouet *et al*., 2018, Redkar *et al*., 2021), and PCR-amplified and cloned 21 of them, prioritizing those that showed allelic variation with other races. Eight CCGs (*i*.*e*. CCG9, CCG12, CCG17, CCG33, CCG35, CCG39, CCG66, CCG80) induced a WRR4A^Oy-0^-dependent HR in *N. tabacum* in a preliminary screen with high *Agrobacterium* concentration (OD_600_ = 0.5) and were further investigated (**Figure S5**). We co-delivered four alleles of WRR4A with the eight candidates at lower *Agrobacterium* concentrations (OD_600_ = 0.4) to test the robustness of their recognition (**Figure 3**). At this concentration, only CCG39 is consistently recognised by WRR4A^Oy-0^ and explains AcEx1 resistance in Oy-0. WRR4A^Col- 0_LONG^ can also recognise CCG39, but WRR4A^Oy-0_SHORT^ cannot. Hence, the C-terminal extension fully explains the acquisition of recognition of CCG39. We also observed a weak recognition of CCG33 by WRR4A^Col-0^, as shown in complementary work (Redkar *et al*., 2021) and which could contribute to the WRR4A-dependent chlorotic response to AcEx1 in Col-0 (**Figure 1**). In addition, WRR4A^Col-0_LONG^ recognises CCG9 and CCG35 (**Figure 3b**). Recognition of CCG9 and CCG35 is not explained solely by the C-terminal extension (as WRR4A^Oy-0^ does not recognise them) or by the core region of the Col-0 allele (as WRR4A^Col-0^ does not recognise them).

WRR4A^Col-0^ can recognise eight CCG effectors from other races of *A. candida* (Redkar *et al*., 2021). We found that WRR4A^Oy-0^ is able to recognise CCG40, CCG104 and CCG28, but not CCG67, CCG79, CCG33, CCG30 and CCG71 (**Figure 3c**). WRR4A^Col-0_LONG^ recognises all the CCGs indistinguishably from WRR4A^Col-0^, indicating no influence of the C-terminal extension on their recognition.

In conclusion, we identified one AcEx1 effector specifically recognised by WRR4A^Oy-0^. The C-terminal extension is required and sufficient for its recognition. We also found that WRR4A^Oy-0^ does not recognise several of the Col-0-recognised CCG from other races.

### WRR4A *alleles carrying a C-terminal extension are associated with AcEx1 resistance*

The NLR repertoire of 64 Arabidopsis accessions has been determined using resistance gene enrichment Sequencing (RenSeq) (Van de Weyer *et al*., 2019). We found 20 susceptible and 5 resistant genotypes that belong to the 64 accessions (**Table S2**). We retrieved *WRR4A* from these 25 accessions (http://ann-nblrrome.tuebingen.mpg.de/apollo/jbrowse/). The read coverage was insufficient to resolve *WRR4A* sequence in Bur-0 (susceptible) and Mt-0 (resistant). *WRR4A* is absent from the *WRR4* cluster in Ws-2, Edi-0 and No-0. Consistently, these accessions are fully susceptible to AcEx1. From the DNA sequence of the 20 other accessions, we predicted the protein sequence, assuming that the splicing sites correspond to those in Col-0 and Oy-0 (**Figure 4**). There are two well-defined clades of *WRR4A* alleles, with a bootstrap value of 100. One clade includes Col-0; the other clade includes Oy-0. The Col-0-like and Oy-0-like clades are also discriminated in a phylogeny based on predicted protein sequences (**Figure S6**). All alleles from the Col-0 clade carry TGA (apart from ULL2-5, TGC, but *WRR4A* is pseudogenised in this accession), while all alleles from Oy-0 clade carry TGC, at the Col-0 stop codon position. Several alleles from both clades, including Bay-0, ULL2-5, Wil-2, Ler-0, Ws-0 and Yo-0, carry an early stop codon (*i*.*e*. upstream of the Col-0 stop codon position), so the resulting proteins are likely not functional. Consistently, all the accessions from the Col-0 clade and all the accessions carrying an early stop codon are susceptible to AcEx1. The only exception is Kn-0, that carries an Oy-0-like allele of *WRR4A* but is susceptible to AcEx1. Otherwise, the presence of an Oy-0-like C-terminal extension associates with resistance.

**Figure 4:**
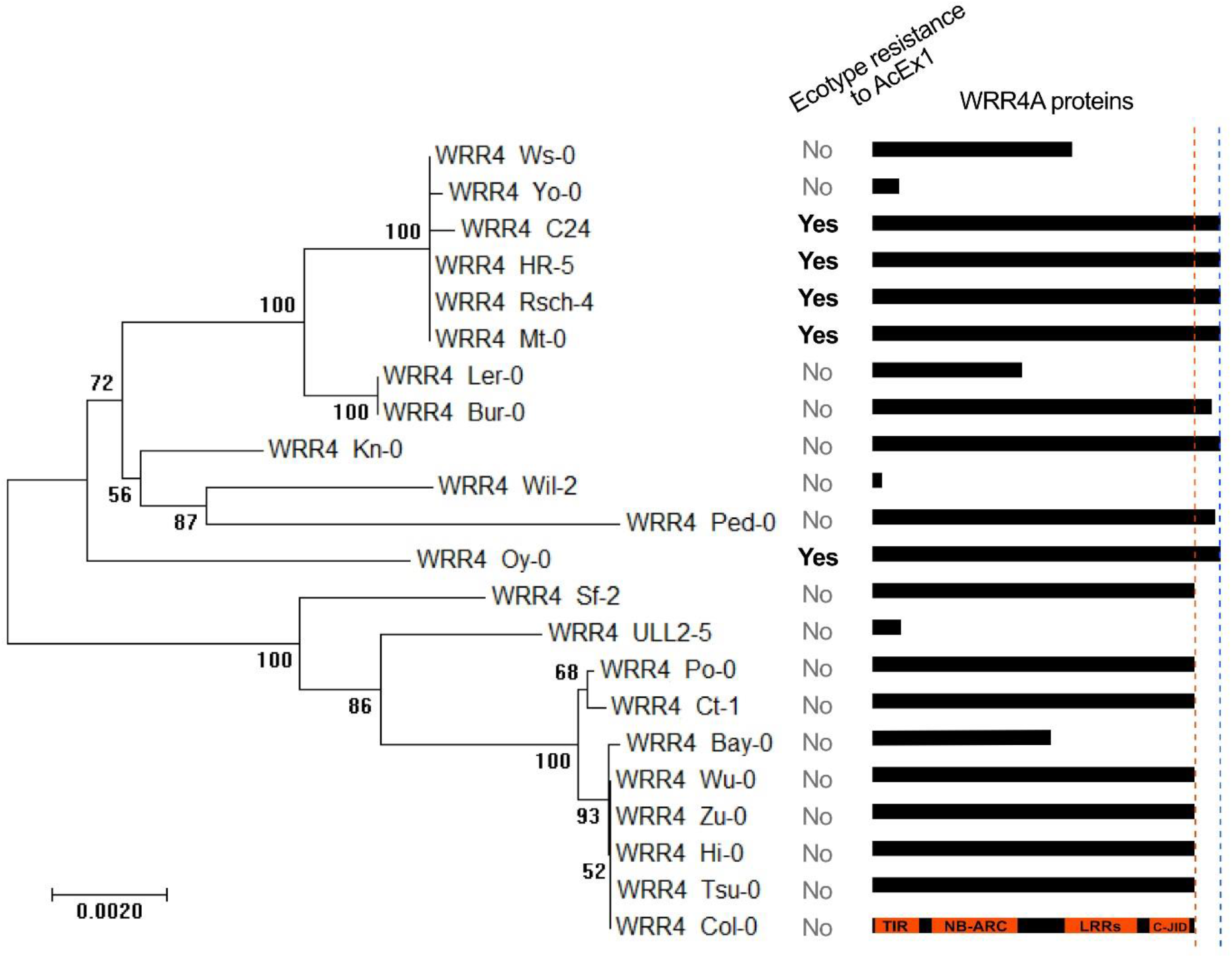
An early stop codon in *WRR4A* is associated with AcEx1 susceptibility. *WRR4A* genomic sequence of 20 Arabidopsis accessions were extracted from http://ann-nblrrome.tuebingen.mpg.de/apollo/jbrowse/ (Van de Weyer *et al*., 2019). Nucleotide sequences corresponding from ATG to TGA of the Oy-0 allele (including introns) were aligned using MUSCLE (software: MEGA7). A phylogenetic tree was generated using the Maximum Likelihood method and a bootstrap (100 replicates) was calculated (software: MEGA7). The tree is drawn to scale, with branch lengths measured in the number of substitutions per site. The resistance / susceptibility phenotypes are indicated. Cartoons on the right represent WRR4A predicted protein, on scale. TIR, NB-ARC, LRR and C-JID are indicated in the Col-0 allele. Dashed orange line represents the Col-0 stop codon. Dashed blue line represents the Oy-0 stop codon.

### AcEx1 resistance can be transferred from Arabidopsis to Camelina

AcEx1 can grow on *Camelina sativ*a **(Figure 5)**. Like Arabidopsis, it can be transformed using the floral dip method (Liu *et al*., 2012). We generated a *WRR4A*^*Oy-0*^-transgenic camelina line. Out of twelve individuals, eight showed complete resistance, three displayed a chlorotic response (likely WRR4A-mediated HR) and one enabled limited pustule formation. All twelve WT camelina control plants showed mild to severe white rust symptoms. This indicates that WRR4A^Oy-0^ can confer resistance to AcEx1 in *C. sativa*.

**Figure 5:**
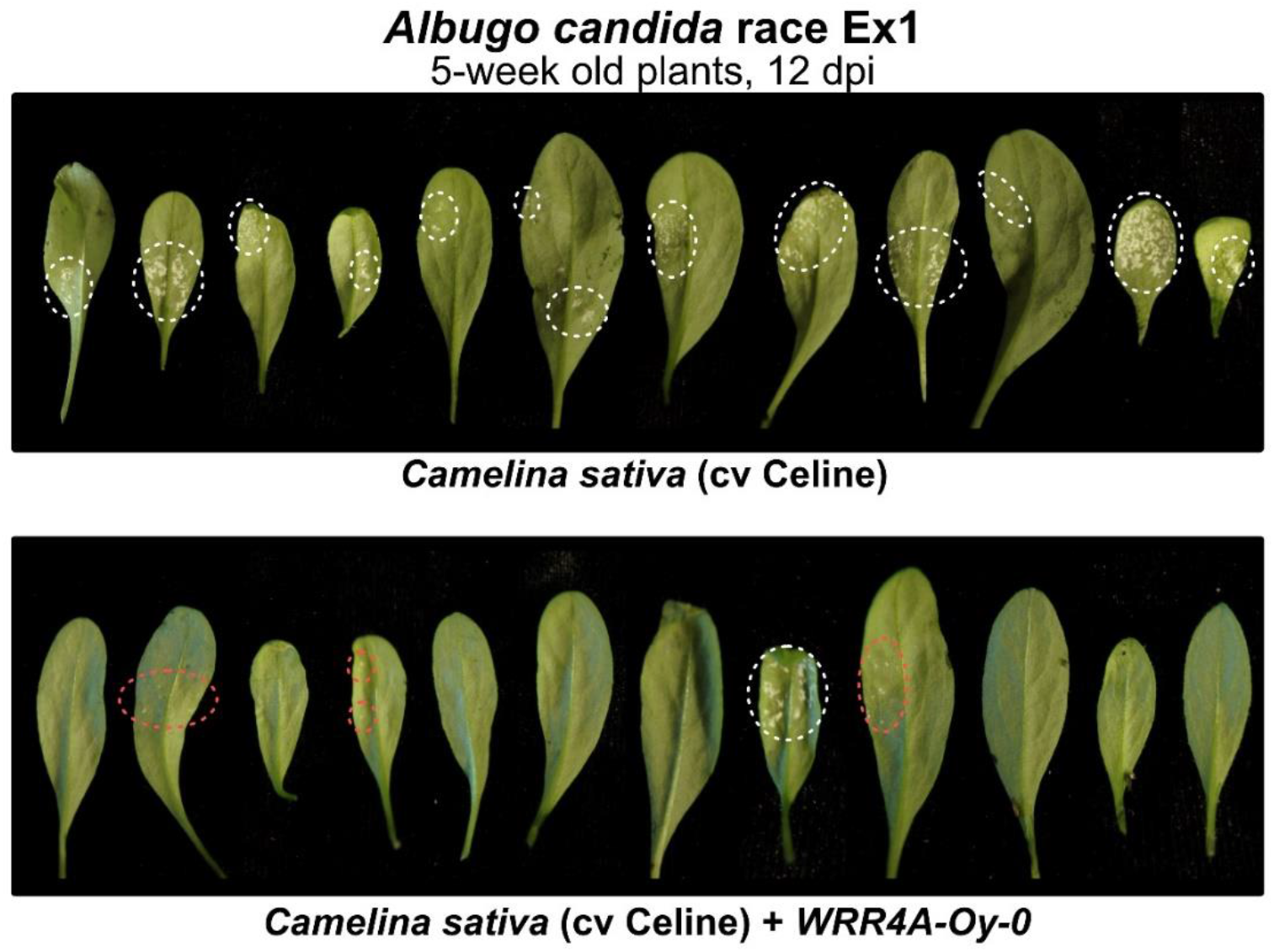
WRR4A confers resistance to AcEx1 in camelina crop. Five-week old camelina (cultivar Celine) plants were sprayed inoculated with AcEx1 race of the white rust oomycete pathogen *A. candida*. Pictures were taken 12 dpi (day post inoculation). **Top row:** Twelve wild-type plants all show mild to severe white rust symptoms. **Bottom row:** twelve lines transformed with *WRR4A*^*Oy-0*^ were tested. One shows mild white rust symptoms, three show local chlorotic response and eight show complete green resistance. White dash line indicates sporulation; red dash line indicates a chlorotic response with no pustule formation.

## Discussion

### Col-0 and HR-5 *WRR4A* alleles recognise effectors from AcEx1

A screen for novel sources of resistance to AcEx1 identified accessions HR-5 and Oy-0 as worthy of further investigation. Positional cloning from Oy-0 and then allele mining in HR-5 showed that this immunity is mediated by alleles of *WRR4A* in HR-5 and Oy-0 with distinct recognition capacities compared to the Col-0 allele. In Oy-0, two additional dominant loci, *WRR11* on chromosome 3 and *WRR15* on chromosome 5 contribute resistance to AcEx1 but the molecular basis of these resistances was not defined. Further investigation on *WRR11* was conducted but did not reveal the causal gene (Castel, 2019, Chapter 3).

WRR4A^Oy-0^ recognizes at least one AcEx1 effector that is not recognized by WRR4A^Col-0^ (**Figure 3**). Conceivably, *WRR4A*^*Oy-0*^ could be combined with *WRR4A*^*Col-0*^ and *WRR4B*^*Col-0*^ to expand the effector recognition spectrum of a stack of *WRR* genes that could be deployed in *juncea* or *C. sativa*.

### WRR4A alleles fall into two clades that can or cannot confer AcEx1 resistance

Analysis of *WRR4A* allele diversity in Arabidopsis revealed WRR4A^Oy-0^-like and WRR4A^Col-0^-like alleles. Since WRR4A^Col-0^-like alleles show near-identity in nucleotide sequence after the premature stop codon to WRR4A^Oy-0^-like alleles, the latter are likely to be ancestral, and the WRR4A^Col-0^-like early stop codon occurred once, in the most recent common ancestor of Sf-2 and Col-0. Other early stop codons, resulting in loss-of-function proteins, occurred randomly in both Oy-0-and Col-0-containing clades. About a third of the investigated accessions contain an early stop codon resulting in a likely non-functional allele (**Figure 4**). The full-length Oy-0-like alleles are associated with resistance to AcEx1, while the Col-0-like alleles are associated with susceptibility (**Figure 3**). The only exception is Kn-0, which displays a full length Oy-0-like allele but is susceptible to AcEx1. Susceptibility in Kn-0 could be explained by SNPs, lack of expression or mis-splicing of *WRR4A*.

### The Col-0 allele C-terminal truncation correlates with gain of recognition for some CCGs and loss of recognition for others, suggesting an evolutionary trade-off

We propose that, in the absence of AcEx1 selection pressure, the Col-0-like early stop codon occurred to provide a new function, along with the loss of AcEx1 effector recognition. This new function enables recognition of more CCGs from other *A. candida* races.

By combining the C-terminal extension on WRR4A^Oy-0^ with the core region of WRR4A in Col-0 (**Figure 3d**), recognition of additional AcEx1 CCGs was enabled. Furthermore, Arabidopsis natural accessions carrying the core region of the Col-0-like allele also lack the C-terminal extension (**Figure 4 and S6**). This could be an example of intramolecular genetic suppression (Kondrashov *et al*., 2002; Schülein *et al*., 2001; Brasseur *et al*., 2001; Davis *et al*., 1999). The combination between the core region of the Col-0 allele with the C-terminal extension may form a hyper-active WRR4A allele with excessive fitness cost for the plant. The early stop codon may have occurred in Col-0 to compensate for hyper-activation of an ancestral WRR4A allele. Hyper-activation of the immune system is deleterious, as shown for example by hybrid incompatibility caused by immune receptors (Wan *et al*., 2021).

Many plant TIR-NLRs show homology in their post-LRR domains (Saucet *et al*., 2020). The recently published structure of Roq1, a TIR-NLR from *N. benthamiana*, reveals a clear-cut C-terminal jelly roll/Ig-like domain (C-JID) after the LRR (Martin *et al*., 2020). It interacts with the Roq1 cognate effector XopQ. Unlike the LRR, the C-JID does not play a role in auto-inhibition. Similarly, in RPP1 (TIR-NLR from Arabidopsis), a C-JID is observed after the LRR (Ma *et al*., 2020). It physically interacts with the cognate effector ATR1 in the active resistosome. We found that WRR4A also contains a PL C-JID. Both WRR4A^Col-0^- and WRR4A^Oy-0^-carry the C-JID, so it does not explain the unique CCG recognition of each allele.

### Arabidopsis *WRR4A* resistance to AcEx1 can be transferred to the crop camelina

*Camelina sativa* was recently engineered to produce LC-PUFAs, an essential component in the feed used in fish farming (Petrie *et al*., 2014). Currently, fish farming uses wild fish-derived fish oil. Fish oil-producing camelina offers a solution to reduce the need for wild fish harvesting, potentially reducing pressure on world marine fish stocks (Betancor *et al*., 2018). There are challenges in delivering products derived from transgenic crops but fish oil-producing crops could reduce the environmental impact of fish farming. White rust causes moderate symptoms on camelina. However, *A. candida* is capable of immunosuppression (Cooper *et al*., 2008). *A. candida*-infected fields constitute a risk for secondary infection of otherwise incompatible pathogens. To safeguard camelina fields against white rust, both chemical and genetic solutions are possible. Genetic resistance offers the advantage of a lower cost for farmers and reduces the need for fungicide release in the environment. Since the first report of white rust on camelina in France in 1945, no genetic resistance has been characterised. All the strains collected on camelina can grow equally irrespective of camelina cultivar (Séguin-Swartz *et al*., 2009). This absence of phenotypic diversity precludes discovery of resistance loci using classic genetic tools. We found that WRR4A^Oy-0^ confers resistance to AcEx1 in camelina (**Figure 5**). Arabidopsis WRR4A resistance is functional in *B. juncea* and *B. oleracea*, suggesting that the mechanism of activation and the downstream signalling of WRR4A is conserved, at least in Brassicaceae.

In conclusion, we found a novel example of post LRR polymorphism within an NLR family, associated with diversified effector recognition spectra. By investigating the diversity of WRRA, we identified an allele that confers white rust resistance in the camelina crop.

## Experimental procedures

### Plant material and growth conditions

*Arabidopsis thaliana* (Arabidopsis) accessions used in this study are Øystese-0 (Oy-0, NASC: N1436), HR-5 (NASC: N76514), Wassilewskija-2 (Ws-2, NASC: N1601) and Columbia (Col-0, NASC: N1092). Col-0_*wrr4a-6* mutant is published (Borhan *et al*., 2008). Seeds were sown directly on compost and plants were grown at 21°C, with 10 hours of light and 14 hours of dark, 75% humidity. For seed collection, 5-weeks old plants were transferred under long-day condition: 21°C, with 16 hours of light and 8 hours of dark, 75% humidity. For *Nicotiana tabacum* (cultivar Petit Gerard) and *Camelina sativa* (cultivar Celine), seeds were sown directly on compost and plants were grown at 21°C, with cycles of 16 hours of light and 8 hours of dark, at 55% humidity.

### Albugo candida Infection assay

For propagation of *Albugo candida*, zoospores from the infected leaf inoculum were suspended in water (∼10^5^ spores/ml) and incubated on ice for 30 min. The spore suspension was then sprayed on plants using a Humbrol® spray gun (∼700 μl/plant) and plants were incubated at 4°C in the dark overnight to promote spore germination. Infected plants were kept under 10-hour light (20 °C) and 14-hour dark (16°C) cycles. Phenotypes were monitored 14 to 21 days after inoculation.

### QTL analysis

QTL mapping of the bipartite F8 Oy-0 x Col-0 population (470 Recombinant Inbreed Lines, RILs, publiclines.versailles.inra.fr/page/27) (Simon *et al*., 2008) was performed on a genetic map of 85 markers across the five linkage groups that accompanied the population using RQTL (Broman *et al*., 2003). Standard interval mapping using a maximum likelihood estimation under a mixture model (Lander and Botstein, 1989) was applied for interval mapping. Analysis revealed two major QTLs: on chromosome 1 and on chromosome 3.

Chromosome 1 QTL is located between 20,384 Mb and 22,181 Mb (**Figure S2**). It includes the TIR-NLR cluster *WRR4* and the CC-NLR cluster *RPP7*. Six RILs (three resistant and three susceptible) recombine within the QTL and were used for fine mapping. We designed a Single Nucleotide Polymorphism (SNP, 21,195 Mb, Fw: TCAGATTGTAACTGATCTCGAAGG, Rv: CCATCAAGCACACTGTATTCC, amplicon contains two SNPs, Oy-0: A and G, Col-0: G and C) and an Amplified Fragment Length Polymorphism (AFLP, 21,691 Mb, Fw: AAGGCAATCAGATTAAGCAGAA, Rv: GCGGGTTTCCTCAGTTGAAG, Oy-0: 389 bp, Col-0: 399 bp) markers between *WRR4* and *RPP7*. Four lines eliminate *RPP7* from the QTL. The only NLR cluster in chromosome 1 QTL is *WRR4*.

Chromosome 3 QTL is located between 17,283 Mb and 19,628 Mb (**Figure S3**). It includes the atypical *resistance*-gene cluster *RPW8*, the CC-NLR *ZAR1* and the paired TIR-NLRs *At3g51560*-*At3g51570*. Six RILs (three resistant and three susceptible) recombine within the QTL and were used for fine mapping. We designed an AFLP (18,016 Mb, Fw: gctacgccactgcatttagc, Rv: CCAATTCCGCAACAGCTTTA, Oy-0: 950 bp, Col-0: 1677 bp) and a Cleaved Amplified Polymorphic Sequence (CAPS, 18,535 Mb, Fw: TCAAGCCTGTTAAGAAGAAGAAGG, Rv: GCCCTCCACAAAGATTCTGAAGTA, enzyme: DdeI, Oy-0: uncleaved, Col-0: cleaved) markers between the QTL border and *RPW8*. We designed a CAPS marker (18,850 Mb, Fw: TCTCGGGGAAAATATGATTAGA, Rv: GGTTGATTTTTATTGTGGTAGTCGT, enzyme: SwaI, Oy-0: cleaved, Col-0: uncleaved) between *RPW8* and *ZAR1*. We designed a SNP (18,937 Mb, Fw: CCACAAGGTCGGAATCTGTAGC, Rv: TGCACAGAAGTAACCCACCAAC, Oy-0: C, Col-0: T) and a CAPS (19,122 Mb, Fw: ACCACCACCTCGATGCATTTC, Rv: CCTTCCCTGCGAAAGACACTC, enzyme: BsrI, Oy-0: uncleaved, Col-0: cleaved) markers between *ZAR1* and the TIR-NLR pair. Three recombinants eliminate the TIR-NLR pair, two eliminate *ZAR1* and one eliminates *RPW8*. None of the Col-0 NLR clusters orthologs are present in the QTL. The gene underlying chromosome three resistance is located between the border of the QTL and RPW8.

### Bulk segregant analysis and RenSeq

We generated an F2 population from a cross between HR-5 (resistant) and Ws-2 (susceptible). We phenol/chloroform extracted DNA from 200 bulked F2 lines fully susceptible to AcEx1. The bulked DNA sample was prepared as an Illumina library and enriched using the c. The sample was sequenced in a pooled MiSeq run (data available on request). Firstly, reads were aligned with BWA mem (Li and Durbin, 2009) to the and SNPs called with Samtools (Li *et al*., 2009). The genome was scanned for regions of high linkage with the next generation mapping tools at http://bar.utoronto.ca/ngm/ (Austin *et al*., 2011). Secondly, the reads were mapped using BWA to the RenSeq PacBio assembly generated for HR-5 (Van de Weyer *et al*., 2019). Highly linked regions were confirmed visually with the integrated genome viewer (Robinson *et al*., 2017).

### Gene cloning

Genes were cloned with the USER method (NEB) following the manufacturer recommendations, with their natural 5’ and 3’ regulatory sequences into LBJJ233-OD (containing a FAST-Red selectable marker, pre-linearized with PacI and Nt. BbvcI restriction enzymes). For overexpression, genes were cloned into LBJJ234-OD (containing a FAST-Red selectable marker and a 35S / Ocs expression cassette, pre-linearized with PacI and Nt. BbvcI restriction enzymes). Primers, template and vectors are indicated in (**Table S3)**. *WRR4A*^*Col-0*^ has been previously published (Cevik *et al*., 2019).

All the plasmids were prepared using a QIAPREP SPIN MINIPREP KIT on *Escherichia coli* DH10B thermo-competent cells selected with appropriate antibiotics. Positive clones (confirmed by size selection on electrophoresis gel and capillary sequencing) were transformed in *Arabidopsis thaliana* via *Agrobacterium tumefaciens* strain GV3101. Transgenic seeds were selected under fluorescent microscope for expression of the FAST-Red selectable marker (Shimada *et al*., 2010).

### CRISPR adenine base editor

An sgRNAs targeting *WRR4A* stop codon in Col-0 (TTCTGAgaagcattcgaaag[nGA]) was assembled by PCR to a sgRNA backbone and 67 bp of the *U6-26* terminator. It was then assembled with the *AtU6-26* promoter in the Golden Gate compatible level 1 *pICH47751*. We designed a mutant allele of a plant codon optimized Cas9 with a potato intron (Addgene: 117515) with D10A (nickase mutant) and R1335V/L1111R/D1135V/G1218R/ E1219F/A1322R/T1337R, to change the PAM recognition from NGG to NG (Nishimasu *et al*., 2018). We assembled this Cas9 (golden gate compatible BpiI: GACA-GCTT) along with a barley codon optimized TadA module (golden gate compatible BpiI: AATG-GCTT) in a level 0 vector *pICH41308*. It was then assembled with the YAO promoter (Addgene: 117513) and the E9 terminator (Addgene: 117519) in a level 1 vector *pICH47811* (with expression in reverse orientation compared to the other level 1 modules). It was then assembled with a FAST-Red selectable marker (Addgene: 117499) and the sgRNA level 1 cassette into a level 2 vector *pICSL4723*, using the end-linker *pICH41766*. Level 0 vector was cloned using BpiI enzyme and spectinomycin resistance. Level 1 vectors were cloned using BsaI enzyme and carbenicillin resistance. Level 2 vectors were cloned using BpiI enzyme and kanamycin resistance. It was expressed via *Agrobacterium tumefaciens* strain GV3101 in Arabidopsis Oy-0. In the first generation after transformation, we did not detect any mutant from 24 independent transformants. It indicates an absence of activity of the construct. It can be explained by the Cas9 mutations that were not tested before on this specific allele nor in combination with TadA.

### *Transient expression in* N. tabacum *leaves*

*A. tumefaciens* strains were streaked on selective media and incubated at 28 °C for 24 hours. A single colony was transferred to liquid LB medium with appropriate antibiotic and incubated at 28 °C for 24 hours in a shaking incubator (200 rotations per minute). The resulting culture was centrifuged at 3000 rotations per minute for 5 minutes and resuspended in infiltration buffer (10 mM MgCl_2_, 10 mM MES, pH 5.6) at OD_600_ = 0.4 (2 ×10^8^ cfu/ml). For co-expression, each bacterial suspension was adjusted to OD_600_ = 0.4 for infiltration. The abaxial surface of 4-weeks old *N. tabacum* were infiltrated with 1 ml needle-less syringe. Cell death was monitored three days after infiltration.

### *Resolution of* WRR4A^Oy*-0*^ *cDNA sequence*

RNA was extracted from three Col-0_*rpw8* and three Col-0 WT individual plants using the RNeasy Plant Mini Kit (QIAgen) and treated with RNase-Free DNase Set (QIAGEN). Reverse transcription was carried out using the SuperScript IV Reverse Transcriptase (ThermoFisher). PCR was conducted using Fw: TCTGATGTCCGCAACCAAAC (in the first exon) and Rv: GTCCTCTTCGGCCATATCTTC (in the last exon) with the Taq Polymerase enzyme (NEB) following the manufacturer protocol. The 2848 nt amplicon sequence, corresponding to the cDNA sequence (*i*.*e*. with already spliced introns) was resolved by capillary sequencing. It indicates that the splicing sites are identical between *WRR4A*^*Oy-0*^ and the splicing sites reported in the database TAIR10 for *WRR4A*^*Col-0*^.

## Acknowledgements

We thank the Gatsby Foundation (UK) for funding to the Jones lab. We thank Mark Youles in TSL Synbio for his excellent support with Golden Gate cloning and for providing modules. This research was supported in part by the NBI Computing infrastructure for Science (CiS) group and Dan MacLean’s group by providing computational infrastructure. B.C., S.F., O.F. and V.C. were supported by Biotechnology and Biological Sciences Research Council (BBSRC) grant BB/L011646/1. A.R. was supported by EMBO LTF (ALTF-842-2015). B.C, S.W and J.D.G.J. were supported in part by ERC Advanced Investigator grant to JDGJ ‘ImmunityByPairDesign’ Project ID 669926.

## Data availability statement

All relevant data can be found within the manuscript and its supporting materials. The sequences of the genomic clones of WRR4A^Oy-0^ and WRR4A^HR-5^ are deposited at NCBI GenBank as MW533532 and MW533533 respectively.

## Author contributions

BC, SF, OF, AR, VC, EH and JJ designed research; BC, SF, OF, AR, SW, VC performed research; BC, SF, OF, AR, VC, EH and JJ analysed data; and BC and JJ wrote the paper. All authors read and approved the final manuscript.

## Conflicts of interest statement

The authors declare that they have no conflict of interests

## Supporting materials

Figure S1: Detailed map of candidate loci in Oy-0

Figure S2: Fine mapping of chromosome 1 QTL

Figure S3: Fine mapping of chromosome 3 QTL

Figure S4: Detailed map of candidate loci in HR-5

Figure S5: Some AcEx1 CCG candidate effectors are recognised by WRR4A^Oy-0^ Figure S6: Phylogeny of WRR4A based on protein sequences

Table S1: Arabidopsis v1 RenSeq bait library (Arbor Bioscience, MI, USA) as described by (Jupe *et al*., 2013)

Table S2: Phenotype of 283 Arabidopsis accessions in response to AcEx1 infection Table S3: Primers used in this study

## Notes

### Competing Interest Statement

The authors have declared no competing interest.

