## Supporting Figures for "Evolutionary trade-offs at the Arabidopsis *WRR4A* resistance locus underpin alternate *Albugo* candida recognition specificities"

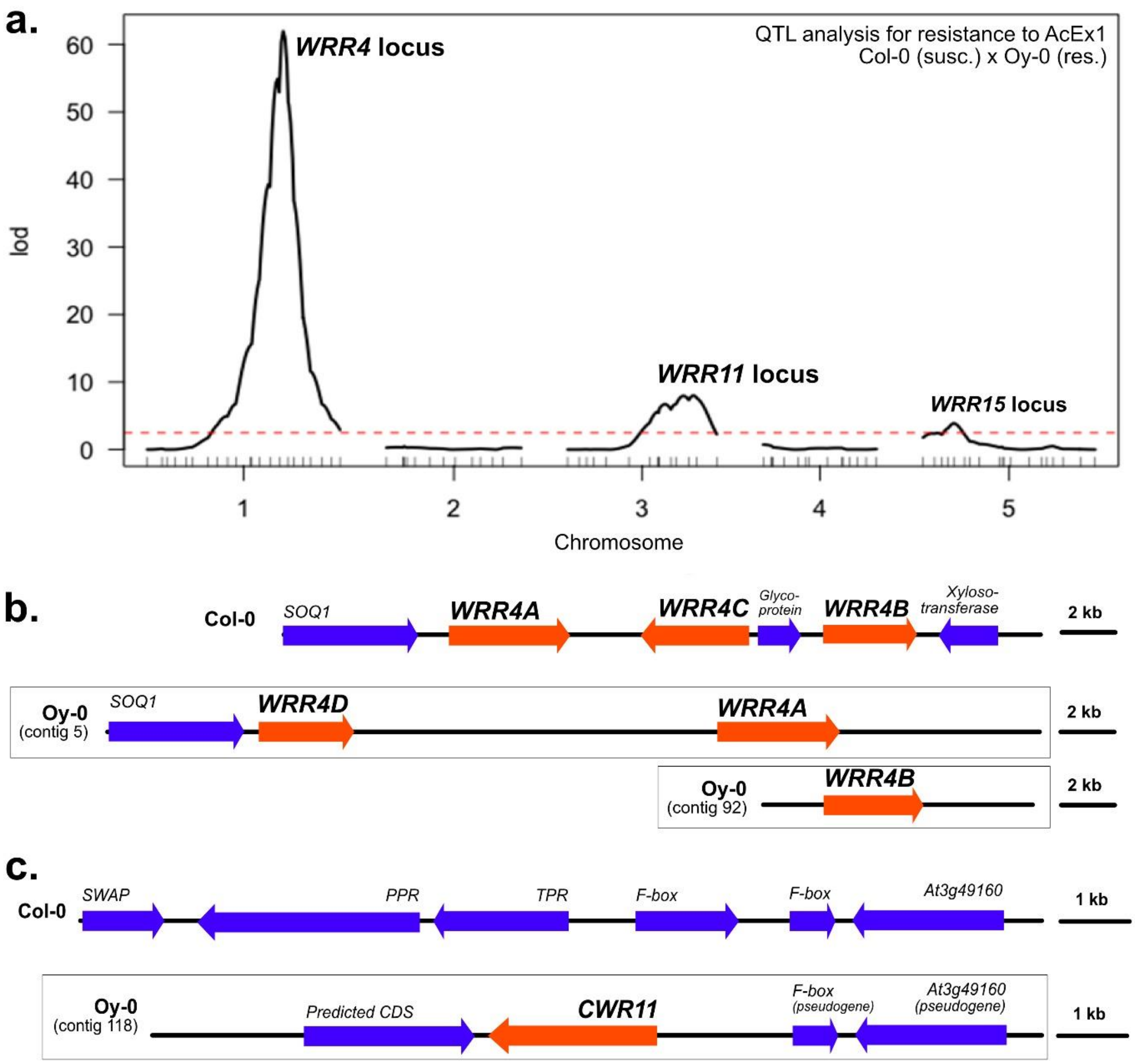


#### Figure S1: Detailed map of candidate loci in Oy-0

**a.** Genome scans using the maximum likelihood algorithm (logarithm of odds (LOD) of 2.5 for both isolates at a 5% confidence interval). Number indicates chromosomes. Three QTLs are over the LOD-score threshold: *WRR4*, *WRR11* and *WRR15*. **b.** *WRR4* locus in Col-0 and two RenSeq contigs containing WRR4 paralogs from Oy-0 (Van de Weyer *et al.*, 2019). **c.** Unique RenSeq contig from Oy-0 sharing identify with the *WRR11* locus. It contains a CC-NLR absent from Col-0. The corresponding locus in Col-0 is displayed on top. Loci are on scale.

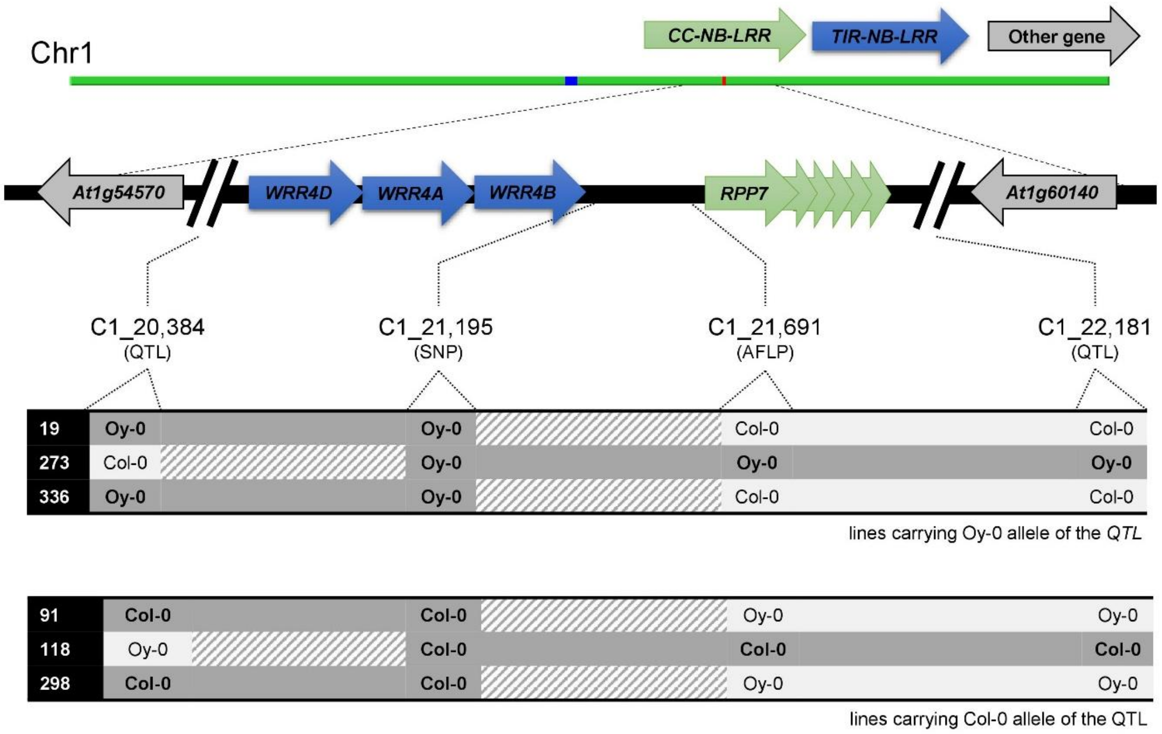


### Figure S2: Fine mapping of chromosome 1 QTL

Six RILs recombines within the Chromosome 1 QTL. Three carry the Oy-0 allele (resistant): 19, 273 and 336. Three carry the Col-0 allele (susceptible): 91, 118, 298. Fine mapping was conducted using two markers between *WRR4* and *RPP7* NLR clusters. Dark grey indicates the region containing the gene. Light grey indicates the region excluding the gene. Dash grey indicates the region containing the Oy-0 / Col-0 recombination site. Numbers indicate the position in Mb on Chromosome 1. QTL: Quantitative Trait Locus (indicates the borders of the QTL before refining). SNP: Single Nucleotide Polymorphism. AFLP: Amplified Fragment Length Polymorphism. Chr1: Chromosome 1. Figure is not on scale.

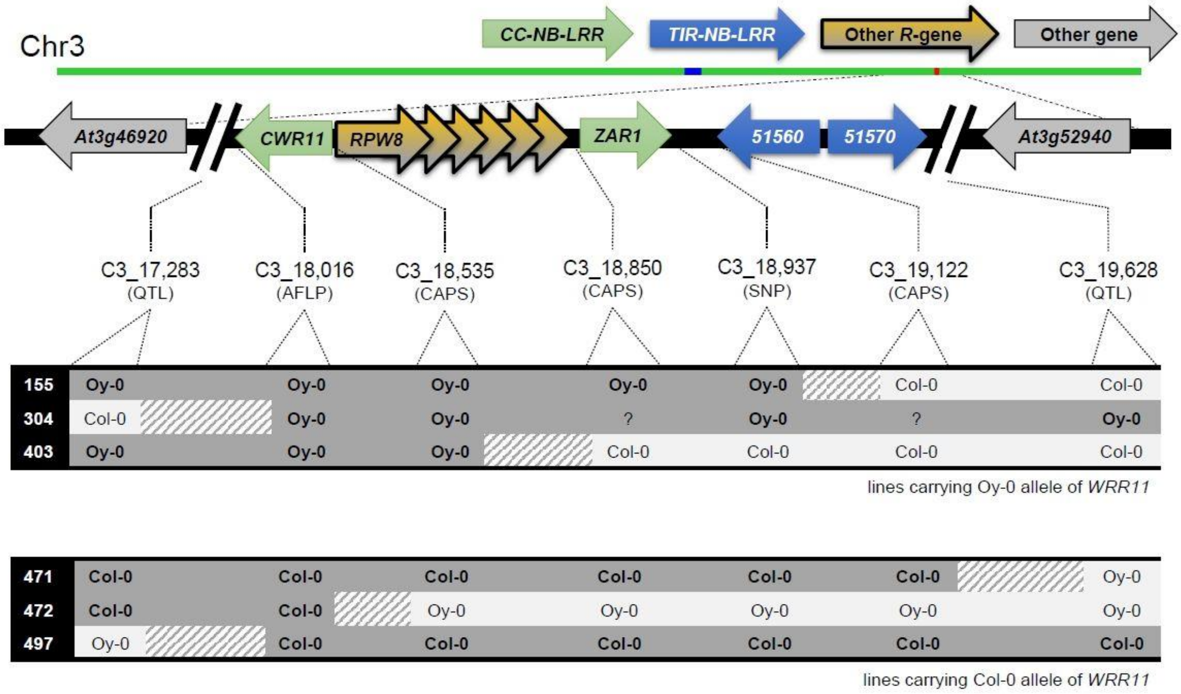


### Figure S3: Fine mapping of chromosome 3 QTL

Six RILs recombines within the Chromosome 3 QTL. Three carry the Oy-0 allele (resistant): 155, 304, 403. Three carry the Col-0 allele (susceptible): 471, 472, 497. Fine mapping was conducted using five markers along the QTL. Dark grey indicates the region containing the gene. Light grey indicates the region excluding the gene. Dash grey indicates the region containing the Oy-0 / Col-0 recombination site. Numbers indicate the position in Mb on Chromosome 3. QTL: Quantitative Trait Locus (indicates the borders of the QTL before refining). SNP: Single Nucleotide Polymorphism. AFLP: Amplified Fragment Length Polymorphism. CAPS: Cleaved Amplified Polymorphic Sequence. Chr3: Chromosome 3. Figure is not on scale.

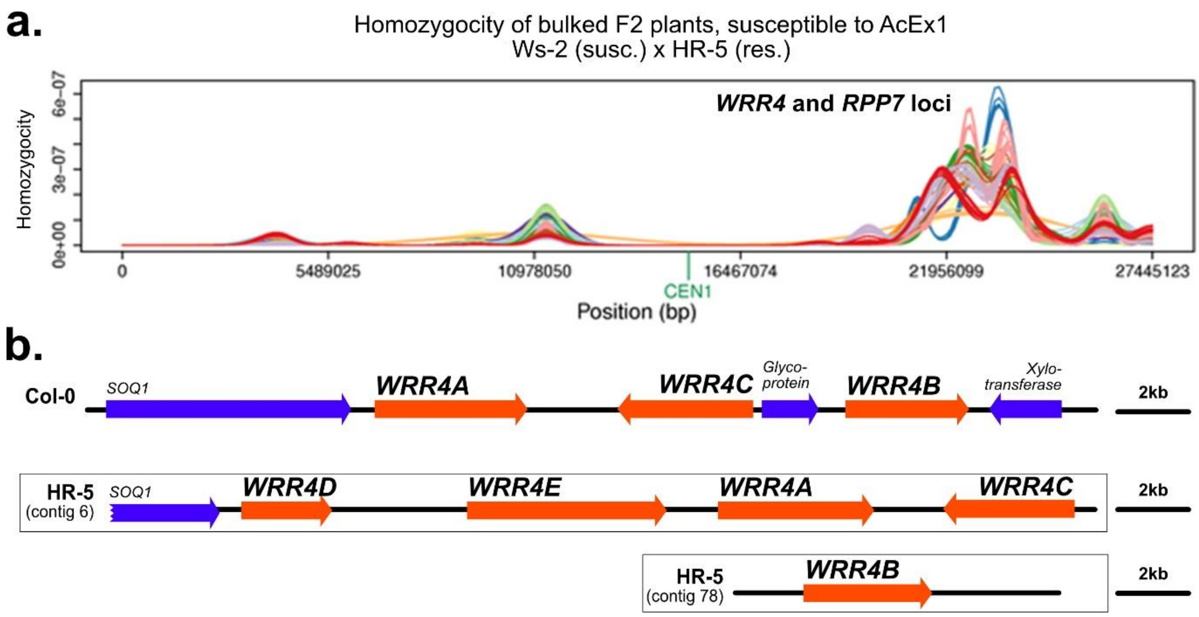


### Figure S4: Detailed map of candidate loci in HR-5

**a.** Homozygosity score of ~100 bulked F2 lines susceptible to AcEx1. Bulked lines were sequence upon *R*-gene enrichment (RenSeq). *WRR4* and *RPP7* cluster present a high degree of homozygosity. This panel was produced using the NGM system where the different coloured bands represent the density of SNPs at different allele frequency levels, used to assess the degree of linkage across the genome. **b.** *WRR4* locus in Col-0 and two RenSeq contigs containing WRR4 paralogs from HR-5 (Van de Weyer *et al.*, 2019).

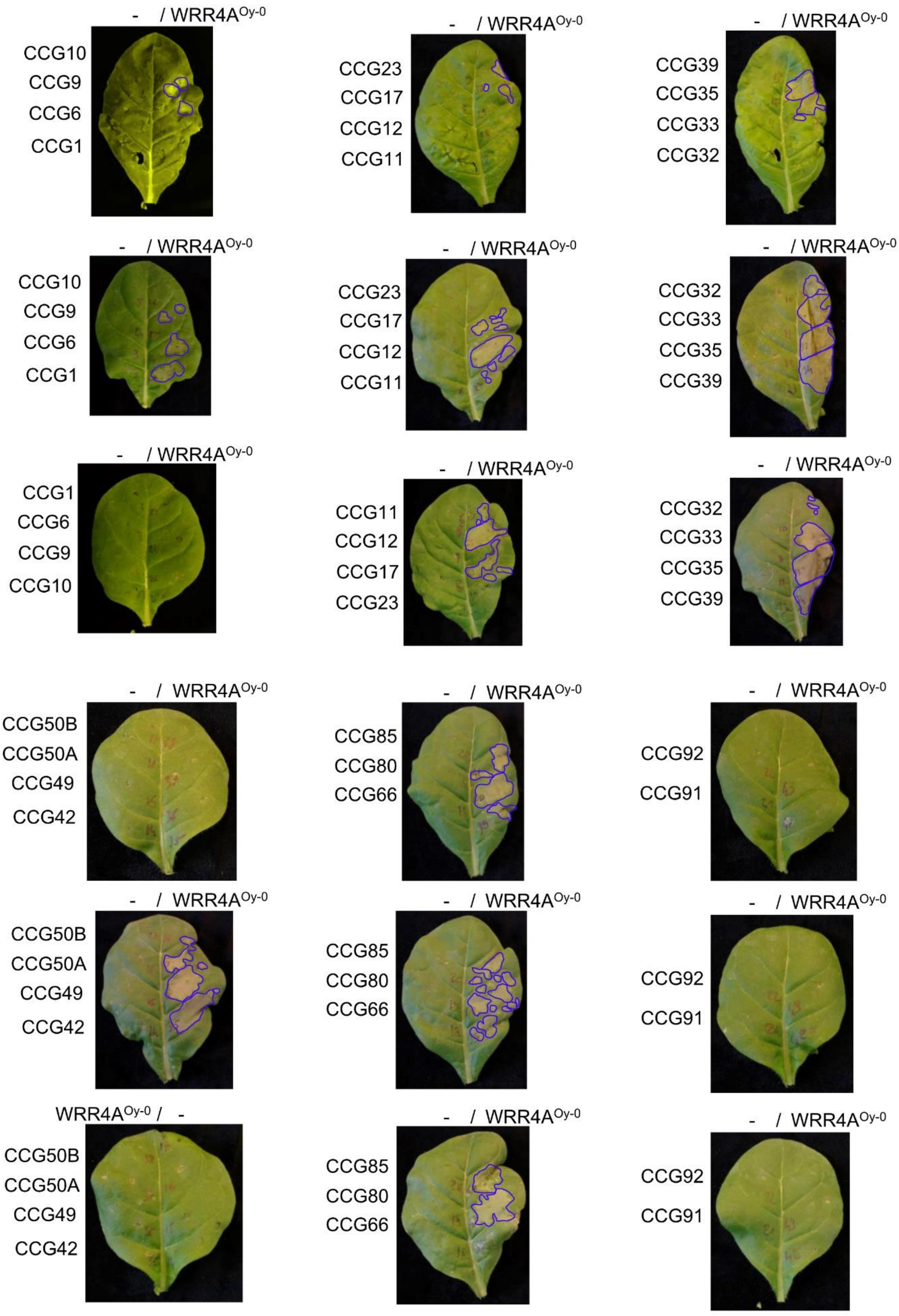


**Figure S5: Some AcEx1 CCG candidate effectors are recognised by WRR4A$^{Oy\text{-}0}$**

4-week old *N. tabacum* plant were infiltrated with *Agrobacterium tumefaciens* strain GV3101 at $OD_{600}$ = 0.5. Genes are expressed under the control of the 35S promoter and the Ocs terminator. Pictures were taken 4 dpi. Blue line indicates HR area, added manually with the software Affinity Designer v1.8.

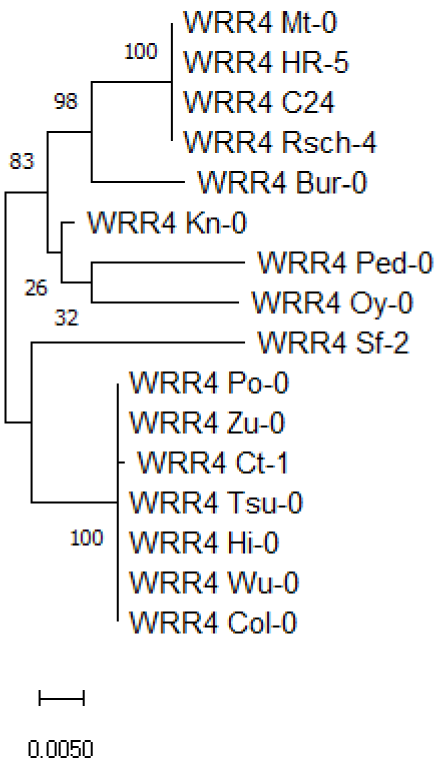


**Figure S6: Phylogeny of WRR4A based on protein sequences**

WRR4A proteins sequence of 16 Arabidopsis accessions were predicted using Augustus (http://bioinf.uni-greifswald.de/augustus/). The C-terminal extension was not used for alignment. Proteins were aligned using MUSCLE (software: MEGA7). A phylogenetic tree was generated using the Maximum Likelihood method and a bootstrap (100 replicates) was calculated (software: MEGA7). The tree is drawn to scale, with branch lengths measured in the number of substitutions per site.
